# Cortical alpha-synuclein preformed fibrils do not affect interval timing in mice

**DOI:** 10.1101/2021.05.10.443485

**Authors:** Qiang Zhang, Hisham Abdelmotilib, Travis Larson, Cameron Keomanivong, Mackenzie Conlon, Georgina M. Aldridge, Nandakumar S. Narayanan

## Abstract

One hallmark feature of Parkinson’s disease (PD) is Lewy body pathology associated with misfolded alpha-synuclein. Previous studies have shown that striatal injection of alpha-synuclein preformed fibrils (PFF) can induce misfolding and aggregation of native alpha-synuclein in a prion-like manner, leading to cell death and motor dysfunction in mouse models. Here, we tested whether alpha-synuclein PFFs injected into the medial prefrontal cortex results in cognitive deficits in mouse models as measured by interval timing, which is reliably disrupted in PD patients and in rodent models. We injected human alpha-synuclein PFF or monomers in the medial prefrontal cortex pre-injected with adeno-associated virus (AAV) overexpressing human alpha-synuclein. Despite notable medial prefrontal cortical synucleinopathy, we did not observe consistent deficits in fixed-interval timing. These results suggest that cortical alpha-synuclein does not reliably disrupt interval timing in rodent models.

**Highlights:** Cortical injection of alpha-synuclein preformed fibrils (PFF) induces diffuse synucleinopathy

Cortical injection of PFFs does not affect interval timing in mice

Medial prefrontal cortical synucleinopathy does not reliably disrupt interval timing

## Introduction

Parkinson’s disease (PD) is a devastating neurodegenerative disorder affecting both movement and cognitive function. PD is characterized by Lewy body pathology, which consists of alpha-synuclein aggregation as well as progressive neurodegeneration of the dopaminergic and cholinergic systems[1-3]. This synucleinopathy can affect cognitive abilities as patients with cognitive impairment often have Lewy bodies and Lewy neurites in the frontal and temporal cortex[4]. Whereas motor symptoms of PD can be managed by dopaminergic medications and deep brain stimulation, cognitive issues related to PD are more difficult to treat[5]. The only FDA-approved medication for symptomatic treatment of cognitive issues in PD is rivastigmine, an acetylcholinesterase inhibitor[1].

One reliable approach to inducing synucleinopathy in rodent models is via preformed fibrils (PFF) of alpha-synuclein. These are a specific alpha-synuclein fibrillar conformation that may facilitate misfolding and aggregation of alpha-synculein monomers. PFFs can act like prions in animal models in that they trigger native alpha-synuclein to fold abnormally and can spread trans-synaptically within neuronal circuits [6-12]. Furthermore, when human alpha-synuclein is overexpressed using adeno-associated virus (AAV) in mice, alpha-synuclein PFF pathology progresses more rapidly[13]. This model can produce reliable cellular and motor deficits when injected in mice[14]. However, studies have reported mixed results regarding cognitive deficits[12, 14].

We investigated cognitive function by injecting human alpha-synuclein PFF into the mouse medial prefrontal cortex bilaterally. To test cognitive function, we used interval timing, which is reliably disrupted in human PD patients and rodent models[5, 15, 16], and depends on the medial prefrontal cortex[17-20]. We used a fixed-interval timing task that requires mice to estimate an interval of 12 seconds and requires working memory for temporal rules and attention to the passage of time[2]. Critically, we have found that fixed-interval timing, is reliably disrupted in rodent PD models[21]. Here, we tested the specific hypothesis that cortical alpha-synuclelin PFFs disrupt fixed-interval timing in mice. Our findings provide insight into cortical alpha-synuclein PFFs and rodent PD models.

## Materials and Methods

### Mice

Fourteen wild-type (WT) C57/BL6J male mice (000664; Jackson Laboratories) were used in this study. All procedures were approved by the Animal Care and Use Committee (#707239) at the University of Iowa, in accordance with the National Institutes of Health Guide for the Care and Use of Laboratory Animals.

### Alpha-synuclein overexpresson

Human alpha-synuclein PFF and monomer solutions were injected into the medial prefrontal cortex of 9-month old mice. At 3 months, animals were pre-injected with a viral vector designed to overexpress either human alpha-synuclein (hSNCA) or control protein (green flourescent protein, or GFP) in targeted regions. Both vectors (CAG-hSNCA-WPRE and CAG-GFP-WPRE) were driven by CAG promoters with recombinant AAV2/6 vectors (serotype 2 genome/serotype 6 capsid), produced and purified by Vector BioLabs (Philadelphia, PA, USA). Titers of AAV were ∼10^13^ GC/ml by qPCR. Alpha-synuclein PFFs and monomers were a gift from the West lab and made according to previously published protocols[7]. Briefly, human alpha-synuclein, encoded in the inducible bacterial expression plasmid pRK172 and purified by maxi-prep (Qiagen), was transformed into chemically-competent BL21 (DE3) Codon Plus cells (Clontech). A single colony was selected for growth in terrific broth (Fisher) supplemented with 100 μg/mL ampicillin (Sigma). Growth was monitored to log-phase protein production induced by isopropyl β-D-1-thiogalactopyranoside (IPTG) for 2 hours at 37°C, and pellets were collected at OD600=0.80. Bacterial pastes were weighed and ∼5 g was combined with 25 mL of lysis buffer (750 mM NaCl, 10 mM Tris (pH 7.6), 1 mM EDTA, 1 mM PMSF, and 1× bacterial protease inhibitor cocktail, RPI). Homogenates were sonicated at 70% power (Fisher Model 500 Dismembrator) for 1 minute 20 seconds (5-second sonication pulses with 25 seconds of off time, on ice), and tubes were placed in boiling water for 15 minutes. After ∼15 minutes on ice, samples were centrifuged for 25 minutes at ∼10,000g for 25 minutes at 4°C, twice. Supernatants were loaded into SnakeSkin dialysis tubing (3.5 kDa MWCO, Fisher) and placed into 4 L of ice-cold buffer (10 mM Tris (pH 7.6) with 50 mM NaCl, 1 mM EDTA, PMSF), stirring overnight. Next, supernatants were centrifuged at 100,000 g for 1 h at 4°C, and concentrated to ∼5 mL using Amicon Ultra-15 3.5 kDa MWCO columns. The concentrate was then passed through a HiLoad 16/600 Superdex Column, 1 ×120 mL (GE Healthcare) with running buffer (10 mM Tris (pH 7.6) with 50 mM NaCl, 1 mM EDTA). Fractions containing alpha-synuclein protein, as assessed by SDS-PAGE and Coomassie stain, were combined and loaded into SnakeSkin dialysis tubing (5 kDa MWCO, Fisher) and placed into 4 L of ice-cold buffer (10 mM Tris (pH 7.6) with 25 mM NaCl, 1 mM EDTA, PMSF), stirring overnight. After dialysis the fractions were loaded at one time into a HiPrep Q HP 16/10 Column, 1 × 20 mL (GE Healthcare) with loading buffer (10 mM Tris (pH 7.6), 25 mM NaCl, l mM EDTA, 1 mM PMSF) and eluted with a gradient application of high-salt buffer (10 mM Tris (pH 7.6), 1 M NaCl, 1 mM EDTA, 1 mM PMSF). Samples containing alpha-synuclein were identified by SDS-PAGE and Coomassie blot, and dialyzed into the same dialysis buffer (excluding PMSF and EDTA) and concentrated as before. Concentration of protein (monomer, undiluted) was determined by BCA assay (Pierce), and purity was assessed by SDS-PAGE and Coomassie blot. Monomer protein diluted to a 5mg/mL solution of human alpha-synuclein protein (buffered with 50 mM Tris (pH 7.5) and 150 mM KCl) in 1.5-mL polypropylene centrifuge tubes were shaken at 700 RPM in an Eppendorf thermomixer for 7 days at 37°C. The solution becomes turbid during the incubation. For sonication of the fibrils, a 1/8-inch probe tip was inserted to a few millimeters from the bottom of the tube, and 30% power (Fisher Scientific Sonic Dismembrator FB120110) was applied for the indicated time (15–240 seconds) in 15-second intervals with 1 sonication pulse and a 1-second wait. Between 15-second intervals, fibrils rested in a metal block at room temperature for 2 minutes to dissipate heat in the solution. Since sonication aerosolizes fibrils, all sonication steps were performed in a biosafety-level-2 cabinet, with the operator wearing disposable wrist guards and double gloves. The sonicated fibrils were then aliquoted and frozen in liquid nitrogen at -80°C, and thawed immediately prior to experiments. For all procedures, monomer alpha-synuclein and fibrillar alpha-synuclein preparations were kept on ice. Cleanup and inactivation of the fibrils on contaminated surfaces was accomplished using a 1%-SDS solution as described[22, 23]. Antibody against tyroxine hydroxylase was purchased from Millipore (AB152). Antibody against phosphorylated alpha-synuclein (p-S129) was purchased from abcam (ab51253). Secondary antibodies (goat anti-rabbit IgG and goat anti-mouse IgG) conjugated to Alexa Fluor 488 and 568, were purchased from Thermo Fisher.

### Mouse fixed-interval timing

At 3 months, mice were trained to perform a fixed interval-timing task with a 12-second interval[21, 24]. Operant chambers (MedAssociates) were equipped with a nosepoke response port, a yellow LED stimulus light (ENV-313W), a pellet dispenser (ENV-203-20), and a house light (ENV-315W). Behavioral chambers were housed in sound-attenuating chambers (MedAssociates). All behavioral responses, including nosepokes and access to pellet receptacles, were recorded with infrared sensors. First, animals learned to make nosepokes to receive rewards (20-mg rodent purified pellets, F0071, Bio-Serv). After fixed-ratio training (FR1), animals were trained in a 12-second fixed-interval timing task in which rewards were delivered for the first nosepoke after a 12-second interval (Figure 2A). The house light was turned on to signal the start of the 12-second interval. Early responses were not rewarded. Responses after 12 seconds resulted in termination of the trial and delivery of the reward. Rewarded nosepokes were signaled by turning off the house light. Each trial was followed by a 30±6-second pseudorandom inter-trial interval that concluded with the house light turning on, signaling the beginning of the next trial. All sessions were 60 minutes long. Mice consumed 1– 1.5 g of sucrose pellets during each behavioral session, and additional food was provided 1–2 hours after each behavioral session in the home cage. Single housing and a 12-hour light/dark cycle were used; all experiments took place during the light cycle. For motivation, mice were maintained at 80%–85% of their baseline body weight during the course of these experiments. Responses were summed into time-response histograms with 1-second bins, 0–18 seconds after the trial start. Plotting was done using the MATLAB function ksdensity.m to estimate the probability-density function of time-response histograms with a bandwidth of 1, normalized to maximum response rate and averaged across animals. We analyzed 10 days of fixed-interval timing performance and quantified behavior by these metrics: 1) the curvature of time-response histograms (1 for a positive curve, and 0 for a flat line[17, 18, 21, 24, 25]); 2) the “start” time of response, calculated using single-trial analysis of trial-by-trial changes between low and high response rates[26, 27] (1–12 seconds); 3) the coefficient of variance (CV) of the start time; and 4) the aggregate number of responses. The median of each of these metrics was calculated over 10 days of behavior and compared between groups by a non-parametric Wilcoxon test.

### Stereotaxic surgical procedures

After 3-month-old mice were trained in the 12-second fixed-interval timing task, they received stereotactical injections of either AAV-hSNCA or AAV-GFP targeting bilateral medial prefrontal cortex, ventral tegmental area (VTA), and basal forebrain. Briefly, mice were anesthetized using ketamine (100 mg/kg) and xylazine (10 mg/kg). A surgical level of anesthesia was maintained, with ketamine supplements (10 mg/kg) given hourly (or as needed) and regular monitoring for stable respiratory rate and absent toe pinch response. Mice were placed in the stereotactic equipment with non-rupturing ear bars. A heating pad was used to prevent hypothermia. Under aseptic surgical conditions, the skull was leveled between bregma and lambda. A Hamilton syringe was lowered to the target areas at the rate of 1 mm/minute, and after a 5-minute delay, AAV was injected slowly at the rate of 0.1 ul/min (0.5 ul per injection site: coordinates from bregma: medial prefrontal cortex: AP: +1.8, ML ± 0.5, DV -1.8; VTA: AP: +3.3, ML ± 1.1, DV -4.6 at 10 degree angled; basal forebrain: AP: -0.7, ML ± 1.8, DV -4.0 and 5.0). Ten minutes after the injection was completed, the syringe was pulled out slowly (2 mm/minute). Following this AAV injection, a second surgery was performed 6 months later to inject either alpha-synulein PFFs or alpha-synuclein monomers into bilateral medial prefrontal cortex (2 ul per injection site (5mg/ml): coordinates from the bregma: medial prefrontal cortex: AP: +1.8, ML ± 0.5, DV -1.8). Mice that were pre-injected with AAV-hSNCA received PFFs, while mice pre-injected with AAV-GFP received monomers.

### Histology

At 15 months when experiments were complete, mice were euthanized by injections of 100 mg/kg ketamine. All mice were intracardially perfused with 4% paraformaldehyde. The brain was removed and post-fixed in paraformaldehyde overnight and immersed in 30% sucrose until the brains sank. Sections (40 µm) were made on a cryostat (Leica) and stored in cryoprotectant (50% PBS, 30% ethylene glycol, 20% glycerol) at -20°C. For immunohistochemistry, briefly, free-floating sections were washed with PBS then PBST (0.03% Triton-X100 in PBS) and incubated in 2% normal goat serum for 1 hour at room temperature, followed by primary antibody incubation at 4°C overnight. After washing with PBST, sections were incubated with secondary antibody for 1 hour. After washing with PBST, then PBS, sections were mounted onto slides with mounting media containing DAPI (Invitrogen P36962). Sections were stained for phosphorylated synuclein at the serine-129 residue (which is linked with enhanced pathogenicity, Abcam, ab51253), tyrosine hydroxylase (Millpore-AB152), and choline-acetyl-transferase (Millipore, AB144P). Images were captured on an Olympus VS120 Microscope. All fluorescent immunohistochemistry images were pseudocolored to represent multiple flourophores.

### Statistics

All procedures were reviewed by the Biomedical Research and Epidemiology Core. Central tendency was reported as median, and variability was reported by intraquartile range. Comparisons were made by non-parametric Wilcoxon rank sum tests. The null hypothesis was tested with Bayes Factor[28].

## Results

### Alpha-synuclein PFFs in medial prefrontal cortex

We injected 9-month-old mice with human alpha-synuclein PFFs into the medial prefrontal cortex. Of note, at 3 months old, mice were pre-injected with AAV-hSNCA into the medial prefrontal cortex, VTA, and basal forebrain. Controls received injections of AAV-GFP in the same locations at 3 months, and alpha-synuclein monomers into medial prefrontal cortex at 9 months. Phosphorylated-alpha-synuclein-(p-syn; S129) positive aggregates with fibrillar morphology were found around the PFF injection site in bilateral medial prefrontal cortex (Figure 1B, C, J), with spread to neighboring prefrontal regions in most mice. We confirmed bilateral pathology at the injection site in all PFF-injected mice. There was no evidence of p-syn-positive cells with fibrillar pathology in control mice (Figure 1D, E). In some mice, we found p-syn-positive cells in the basal forebrain where we overexpressed human alpha-synuclein though AAV, but there were no soma with clear fibrillar pathology (Figure 1L). In the VTA and substantia nigra, we found scattered cells with fibrillar p-synuclein (0–2 postive soma per section) (Figure 1F). Given the lack of significant fibril spread to this region, we stained for tyrosine hydroxylase in two mice per group. As expected, we saw no gross differences in TH staining in these mice (Figure 1G, H), in line with the lack of significant fibrillar spread to this region. Unexpectedly, we found moderate amounts of p-synuclein in a fibrillar pattern in soma and neurites in the temporal cortex, including the entorhinal cortex (Figure 1I). These regions were not preinjected with virus coding for human alpha-synuclein. We also found soma with fibrillar staining in the striatum, though the density was more limited compared with the medial frontal cortex (Figure 1K).

**Figure 1:**
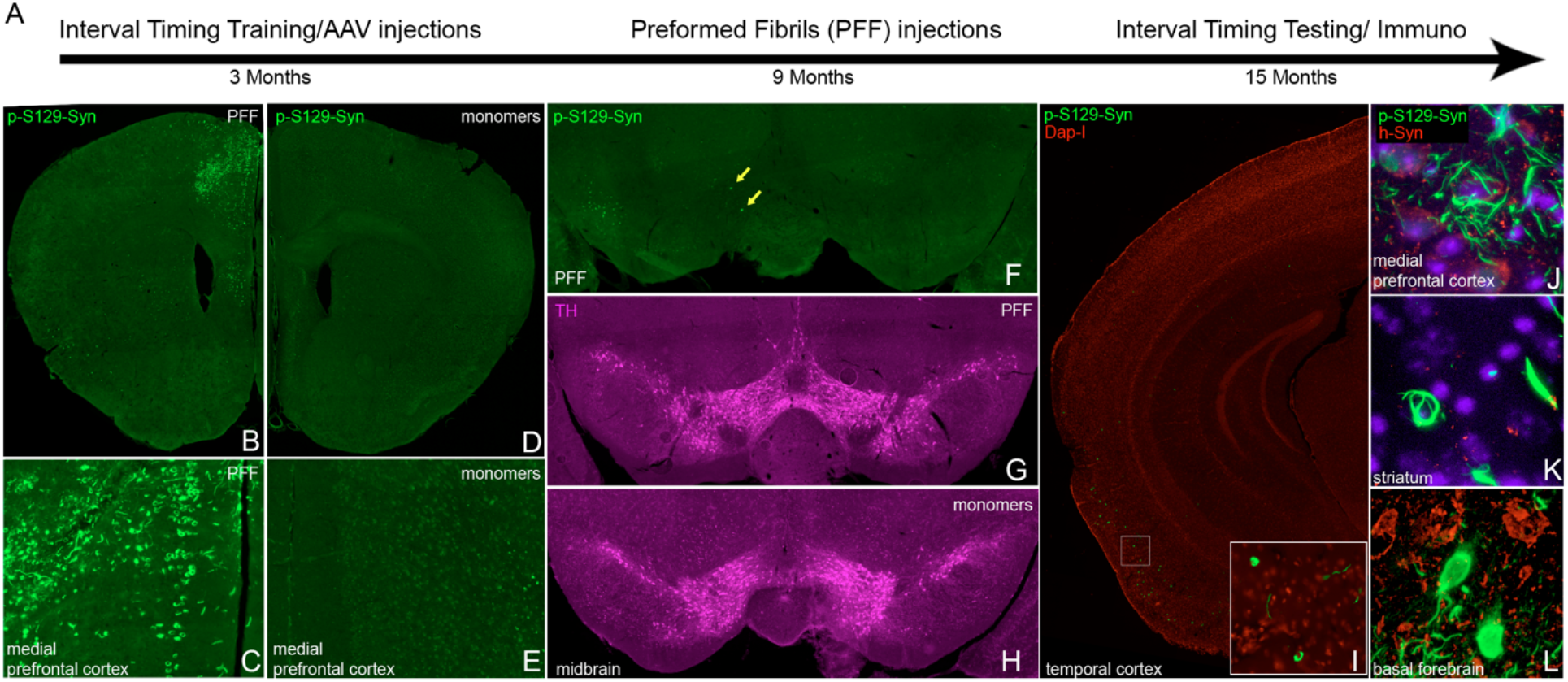
Human alpha-synuclein preformed fibrils caused extensive local pathology after injections into bilateral medial prefrontal cortex. A) Timeline of the experiment: 3-month-old wild type mice were trained for a 12-second fixed-interval timing task prior to receiving AAV injections for overexpression of human alpha-synuclein in bilateral medial prefrontal cortex, basal forebrain, and ventral tegmental area (VTA). Control mice received AAV-GFP injections to overexpress green fluorescence protein (GFP). After 6 months, 9-month-old mice received medial prefrontal cortical injections of human alpha-synuclein preformed fibrils (PFF) (AAV-hSNCA group) or human alpha-synuclein monomers (AAV-GFP group). After another 6 months, fixed-interval timing testing was performed and animals were euthanized for immunohistochemistry. B-C) Extensive phospho-synuclein (p-syn) pathology was noted in the medial prefrontal cortex at the level of the injection site. D-E) P-Synuclein pathology was not detected in the control group. Fluorescent images were pseudocolored and intensity-matched for background fluorescence. Differing secondary antibodies were used for groups due to control group injection of GFP. F) Scattered p-syn+ soma (yellow arrows) with fibrillar pathology were rare in VTA/substantia nigra. G-H) Tyrosine hydroxylase (TH) staining in PFF-(G) and monomer-(H) injected animals. I) p-syn+ soma with fibrillar pattern were found in moderate densities in temporal cortex. J-L) confocal microscopy images confirmed fibrillar p-synuclein pathology in the medial prefrontal cortex (J), the injection site of PFF, and the striatum (K), which was not a site of viral overexpression. The basal forebrain (L), a site of viral overexpression, showed p-syn+ soma with diffuse, rather than fibrillar staining.

### Cortical alpha-synuclein PFFs do not affect interval timing

Because injection of PFFs into the striatum can disrupt motor behaviors[29], we tested the hypothesis that cortical PFFs can disrupt cognitive behaviors. We turned to interval timing, which is reliably disrupted in human PD patients[5, 30], correlated with cognitive dysfunction in these patients, and disrupted in rodent PD models that disrupt the medial prefrontal cortex[21, 31]. One reliable metric that is disrupted by dopaminergic manipulations is the *curvature* of time-response histograms, which uses the cumulative distribution to measure deviations from a flat line, ranging from -1 to 1, with 0 being a flat line[17]. Strikingly, we found that despite observing clear synucleinopathy in the medial prefrontal cortex, there was no consistent deficit in the curvative of time-response histograms (Median(IQR): Control 0.18(0.17-0.22), PFF 0.18(0.14-0.24); p>0.99; Bayes Factor 2.8), start times (Control 5.5(5.1-6.0), PFF 6.0(5.3-6.0); p=0.7; Bayes Factor 2.5), start time CVs (Control 0.32(0.30-0.33), PFF 0.34(0.33-0.36); p=0.10; Bayes Factor 1.4), or nosepokes (Control 190(96-272), PFF 148(92-192); p=0.71; Bayes Factor 2.0) (Figure 2). There was no reliable difference in any metric of timing that we could measure. These data provide evidence that fibrillar synuclein in the medial prefrontal cortex does not reliably impair fixed-interval timing in mice.

**Figure 2:**
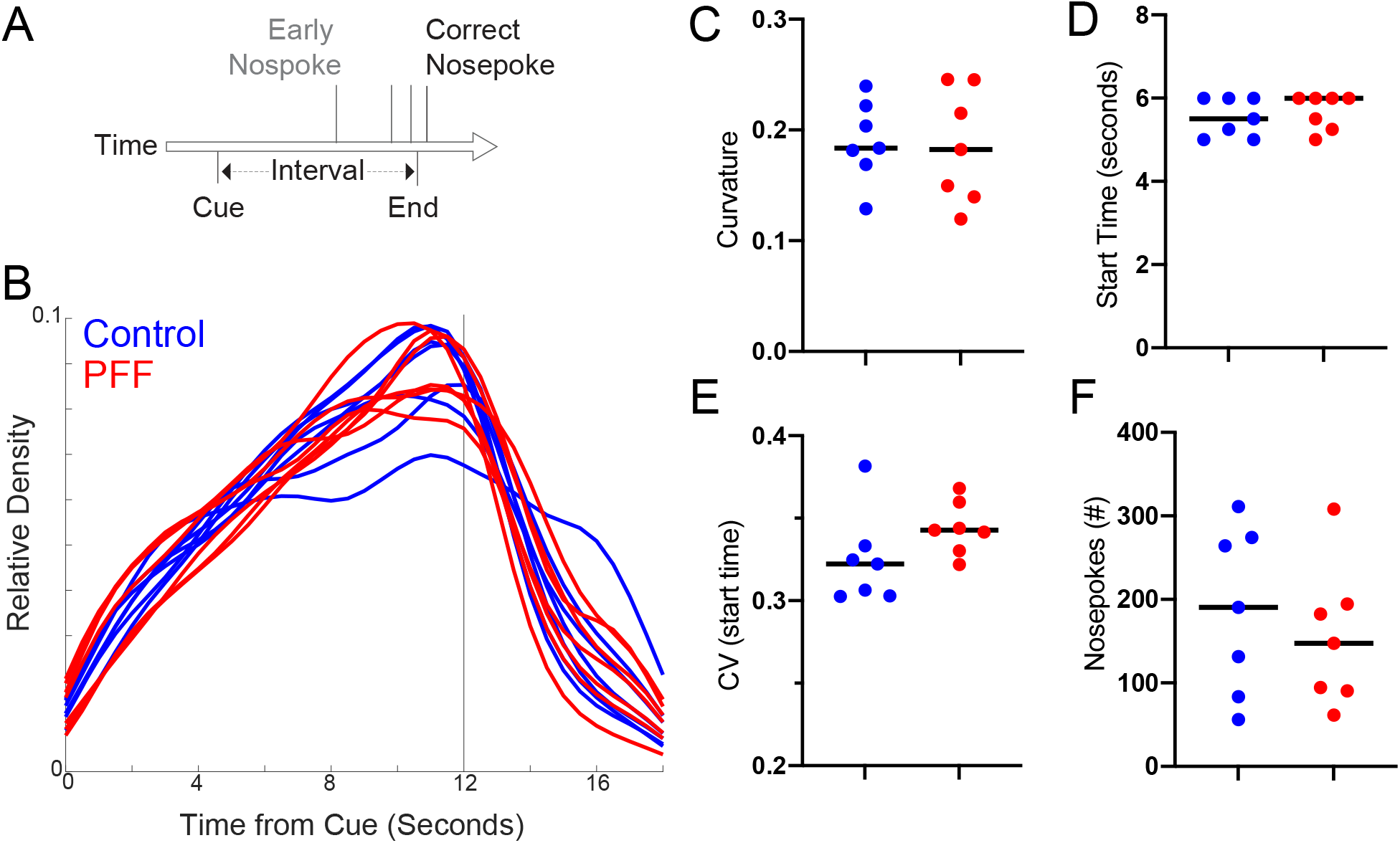
Fixed-interval timing was not affected by cortical PFF-induced synucleinopathy. A) We trained 3-month-old mice to perform a 12-second fixed-interval timing assay and then injected animals with cortical PFFs. The mice estimated a 12-second interval by making a nosepoke response; the first nosepoke after 12 seconds was rewarded. B-F) We found no consistent differences in fixed-interval timing between control (blue) and PFF (red) animals in time-response histograms (B), or in C) the curvature of time-response histograms, D) the time animals began nosepoking, as measured by the start time from single-trial analysis, E) the coefficient of variation (CV) of start times from single trial analysis averaged over 10 days, or F) the total number of responses. Data from PFF (n=7) and control (n=7) mice, 6 months after injection.

## Discussion

We tested the hypothesis that medial prefrontal cortical alpha-synuclein PFFs affect cognitive function in rodent models. Despite evident synucleinopathy in this region, we did not find consistent evidence of cognitive deficits in a fixed-interval timing task that is reliably impaired in rodent PD models. We note that interval timing is reliably impaired in PD. Recent work from our group found that medicated PD patients have strongly increased variance of responses during interval timing. Furthermore, past work from our group and others has consistently shown that disrupting midbrain dopamine[17, 21, 32, 33], cortical dopamine[30, 34], or striatal dopamine reliably impairs interval timing[35, 36]. Given these data, it is interesting that cortical PFFs have no measurable timing impairments.

There are three possibilities for our results. First, cortical cells may be less vulnerable to alpha-synuclein fibrils compared with other brain regions. Blumenstock et al. showed abnormal morphology in cortical pyramidal cells following PFF injections in the striatum after five months[37]. In that study, fibrils were derived from mouse alpha-synuclein, which has been shown in some cases to cause more spread and pathology compared to human fibrils used in our study[7]. Although we did see extensive alpha-synuclein pathology in the medial prefrontal cortex, it is possible that more widespread pathology in the cortex would be required to see a deficit in this task. It is also possible that alpha-synuclein perturbations in the cortex act distinctly from subcortical alpha-synuclein[38]. Second, interval-timing deficits may depend on pathology in subcortical projection neurons, such as dopaminergic projection neurons in the VTA, as we have shown previously[17, 21, 39]. Although we found spread to subcortical regions such as the striatum, the most dense fibrillar pathology was in the medial prefrontal cortex (Figure 1). In this model, there was almost no fibrillar staining of synuclein seen in the VTA and substantia nigra. Although it is possible that the apparent lack of spread could be secondary to cell loss, this is unlikely, given the lack of behavioral deficits. Finally, it is possible that cortical networks compensated for synucleinopathy. Future studies might unravel these possibilities in describing the role of cortical alpha-synuclein in cognitive function.

Our study has limitations. First, our control animals received both AAV-GFP and human alpha-synuclein monomer injections, with no separate vehicle-only control. However, our curvature metrics from the control mice closely match our prior work, suggesting these manipulations did not alter behavior[21, 24]. Second, we did not extensively evaluate dopaminergic cell loss in this mouse model. As there was no behavioral difference between groups, we chose not to quantitatively evaluate dopaminergic cell loss in the VTA or substantia nigra. Qualitatively, there was no evidence of a substantial decrease in TH staining in the sections we examined, consistent with our hypothesis that dopaminergic cell loss is key to interval-timing deficits in mouse models of PD. Third, we only used one measure of timing (12-second fixed-interval timing). Given the complexity and multiple surgeries required for this study, we chose to use the simplest measure of interval-timing. It is possible other aspects of cognition or timing could be influenced by cortical alpha-synuclein pathology.

Despite these limitations, our data provide evidence that injection of human alpha-synuclein PFFs into the medial prefrontal cortex does not reliably disrupt interval timing in rodent models, which may guide the choice of injection site and fibril forms in future investigations using alpha-synuclein models of synucleinopathy.

## Author contributions

QZ and NN conceived and designed the study, GA, HA, TL, CK, MC and QZ carried out the experiments, GA, HA, QZ and NN performed data analysis, QZ, HA and NN wrote the manuscript.

## Acknowledgements

QZ is supported by NIH/NINDS R25 NS079173, the NIH/NINDS NeuroNext Fellowship, and the physician scientist training program at University of Iowa. QZ is a trainee of the University of Iowa Clinical Neuroscientist Training Program (CNS-TP). NN is supported by NIH R01 NS100849-A1. GMA, NIH K08 NS109287, and the Iowa Neuroscience Institute.

